# Easymap: a user-friendly software package for rapid mapping by sequencing of point mutations and large insertions

**DOI:** 10.1101/2021.01.06.425624

**Authors:** Samuel Daniel Lup, David Wilson-Sánchez, Sergio Andreu-Sánchez, José Luis Micol

**Affiliations:** Instituto de Bioingeniería, Universidad Miguel Hernández, Campus de Elche, 03202 Elche, Spain

**Keywords:** mapping-by-sequencing software, candidate mutations, forward genetics, NGS

## Abstract

Mapping-by-sequencing strategies combine next-generation sequencing (NGS) with classical linkage analysis, allowing rapid identification of the causal mutations of the phenotypes exhibited by mutants isolated in a genetic screen. Computer programs that analyze NGS data obtained from a mapping population of individuals derived from a mutant of interest in order to identify a causal mutation are available; however, the installation and usage of such programs requires bioinformatic skills, modifying or combining pieces of existing software, or purchasing licenses. To ease this process, we developed Easymap, an open-source program that simplifies the data analysis workflows from raw NGS reads to candidate mutations. Easymap can perform bulked segregant mapping of point mutations induced by ethyl methanesulfonate (EMS) with DNA-seq or RNA-seq datasets, as well as tagged-sequence mapping for large insertions, such as transposons or T-DNAs. The mapping analyses implemented in Easymap have been validated with experimental and simulated datasets from different plant and animal model species. Easymap was designed to be accessible to all users regardless of their bioinformatics skills by implementing a user-friendly graphical interface, a simple universal installation script, and detailed mapping reports, including informative images and complementary data for assessment of the mapping results.

**One sentence summary:** Easymap is a versatile user-friendly software tool that facilitates mapping-by-sequencing of large insertions and point mutations in plant and animal genomes

## INTRODUCTION

Forward genetic screens consist of random mutagenesis followed by the isolation of mutants exhibiting a phenotype of interest, and genetic analysis of these mutants to identify the mutations that cause their phenotypes. Two commonly used mutagenesis strategies are the induction of G→A substitutions using the chemical mutagen ethyl methanesulfonate (EMS) (Neuffer and Ficsor, 1963; James and Dooner, 1990; Jansen et al., 1997) and the disruption of genes by insertional mutagens such as transposons or T-DNAs (Cooley et al., 1988; Alonso et al., 2003; Frøkjær-Jensen et al., 2014). Linkage analysis of molecular markers in segregant mapping populations is the classically preferred approach to map the point mutations induced by a chemical mutagen, carried by mutants isolated in a genetic screen (Michelmore et al., 1991; Ponce et al., 1999). By contrast, localization of insertional mutations has relied on methods to capture the genomic sequences present at their flanks (Gasch et al., 1992; Medford et al., 1992; Liu et al., 1995; O’Malley et al., 2007).

The preliminary identification and subsequent validation of the mutations that cause a phenotype of interest is the most laborious and time-consuming step of a forward genetic screen. Next-generation sequencing (NGS) of DNA has facilitated and revitalized such approaches, through the so-called mapping-by-sequencing methods, which combine NGS with linkage analysis (Schneeberger and Weigel, 2011; Hartwig et al., 2012; James et al., 2013; Candela et al., 2015). Mapping-by-sequencing approaches for the identification of causal mutations are much faster than previous methods but can be hampered by the lack of computing resources and/or accessible software. Currently available programs for mapping-by-sequencing data analysis suffer from one or several of the following issues: they require the purchase of licenses (Smith, 2015); they require a certain level of bioinformatics skills to use (Abe et al., 2012; Fekih et al., 2013; Jiang et al., 2015; Sun and Schneeberger, 2015; Ecovoiu et al., 2016; Wachsman et al., 2017) (see also https://sourceforge.net/projects/mimodd/); they only do a part of the computing tasks required for a mapping-by-sequencing experiment (Li et al., 2009; Langmead and Salzberg, 2012; Hill et al., 2013); they are designed for a specific type of mutation or mapping strategy (Gonzalez et al., 2013; Ewing, 2015; Hénaff et al., 2015; Solaimanpour et al., 2015; Sun and Schneeberger, 2015; Wachsman et al., 2017; Klein et al., 2018; Javorka et al., 2019); they are hosted at a public server but usage is limited (Gonzalez et al., 2013; Afgan et al., 2018); or they can no longer be accessed or used (Minevich et al., 2012; Pulido-Tamayo et al., 2016).

Here we describe Easymap, an accessible graphical-interface program that analyzes NGS data from mapping populations derived through a variety of experimental designs from either insertional or EMS-induced mutants. This software package avoids the above-mentioned issues, enabling mapping-by-sequencing experiments to be conducted by researchers with minimal bioinformatics experience. Easymap features a web-based graphic interface, a simple installation script, robust mapping analyses for several experimental designs, and thorough user-oriented mapping reports.

## RESULTS

### Mapping strategies for which Easymap can be used

Easymap offers the user two alternative workflows, to map point and insertional (Workflows 1 and 2; see below), as well as a set of complementary tasks common to all analyses. Figure 1 offers an overview of both mapping strategies, from the initial selection of the mutants of interest to the output that Easymap generates. To ease both mapping scenarios, Easymap automates the whole data analysis process requiring no user intervention with alignment, variant-calling or filtering parameters.

**Figure 1.**
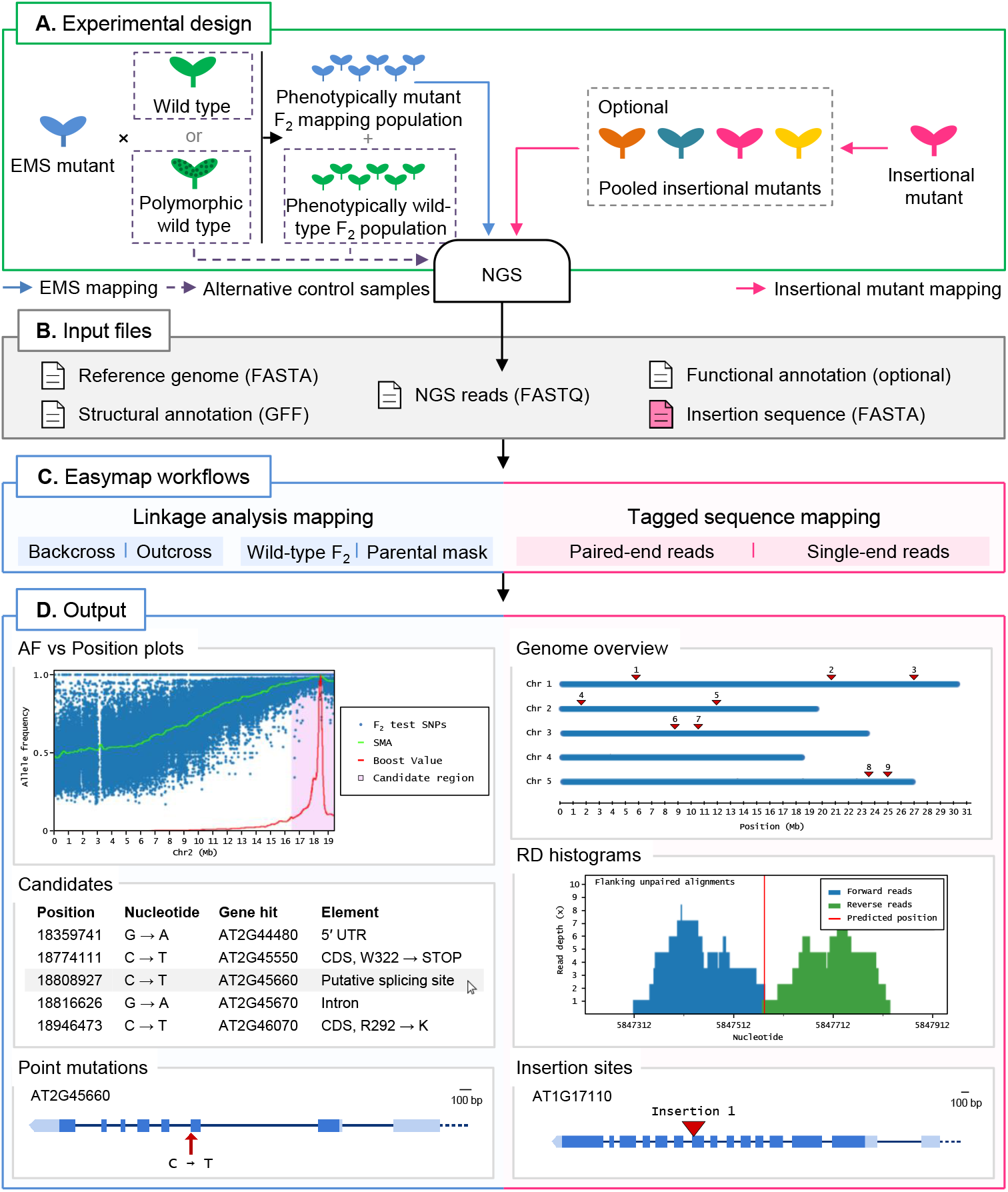
Overview of a mapping-by-sequencing experiment with Easymap in Arabidopsis. (A) Experimental design. For EMS-induced mutants, an outcross or backcross is first performed. The F_1_ plants derived from the cross are selfed, and the resulting F_2_ is screened for the mutant phenotype to create a phenotypically mutant mapping population. A control sample is required for the mapping analysis, which can be either one of the parental individuals crossed or, alternatively, a pool of phenotypically wiid-type F_2_ individuals. For mapping of large insertions, the DNA of different insertional mutant lines can be sequenced individually or pooled, and no control sample is required. (B) Input files. Easymap takes NGS paired-end or single-end short reads as input. The remaining mandatory input files are available on public databases for each model species. (C) Easymap workflows. The user selects the experimental design used for mutation mapping from a variety of options for both EMS mutation mapping (backcross and outcross strategies, alternative control samples) and tagged-sequence mapping (paired-end and single-end reads). (D) Output. Easymap produces comprehensive mapping reports with organized tabular data to ease interpretation of the results. As an example of EMS-induced mutations, data from the Arabidopsis *suppressor of overexpression of CONSTANS 1-2 (soc1-2)* mutant (Sun and Schneeberger, 2015) was used for this figure. Allele frequency (AF) versus Position plots are drawn for each chromosome containing the polymorphisms used for the analysis. A candidate region is highlighted in pink; all putative EMS-type mutations contained in this region are regarded as candidates, and their position and relevant information is presented in a table, such as the gene affected by the mutation. For each gene affected by a candidate mutation, a gene plot is made in which the position of the mutation is shown, followed by further information (genotyping primers, flanking sequences, functional annotation, etc.). As an example of large insertion mapping, the figure includes data from an unpublished mapping experiment made in our laboratory (see Table 3). A genomic overview is drawn showing the positions of the insertions found. Read depth (RD) histograms are generated for each read cluster pointing to an insertion site showing the information supporting the insertion. Finally, a gene plot is made for each gene interrupted by an insertion.

#### Workflow 1: Linkage analysis mapping

Mapping of causal EMS-induced mutations is typically achieved by linkage analysis in a bulked segregant population. The user must obtain a mapping population by phenotyping the M_2_ offspring of an M_2_ individual or the F_2_ offspring of a backcross (in which an M_2_ mutant is crossed to an individual genetically identical to the parent subjected to mutagenesis) or an outcross (in which an M_2_ mutant is crossed to an individual unrelated and genetically polymorphic to the parent that was subjected to mutagenesis).

Easymap includes the splice-aware aligner HISAT2 (Kim et al., 2019), which is three times faster than commonly used aligners in default conditions with no impact on memory usage or sensitivity. The implementation of HISAT2 allows the user to input RNA-seq and DNA-seq reads indistinctly for point mutation mapping. Easymap requires NGS reads from test and control samples. The test sample consists of NGS reads obtained from a population of individuals exhibiting the mutant phenotype of interest—hence homozygous for a recessive mutation that causes that phenotype. The control sample can be pooled M_2_ or F_2_ phenotypically wild-type individuals, or individuals genetically identical to the parent that was subjected to mutagenesis, or the strain to which the mutant is outcrossed. A minimum coverage of 25× is recommended for each sample, although since coverage directly correlates to the reliability of the results, higher coverages are encouraged. Table 1 and the Easymap Documentation (File S1) describe the different experimental designs supported by Easymap, four of which are detailed in Figure 2. Easymap first calls the single-nucleotide polymorphisms (SNPs) between the reads obtained by the user and the reference sequence for the genome of the species under study, and identifies high-confidence SNPs that are informative for mapping. The allele frequencies of these biallelic markers are then analyzed to predict the phenotypically selected genomic position, which defines the center of a candidate interval predicted to contain the causal mutation. The SNPs in the candidate region are then analyzed and reported as candidates to be the mutation causing the phenotype under study. The Easymap documentation (File S1) contains more detailed information about the SNP selection and mapping algorithms implemented.

**Figure 2.**
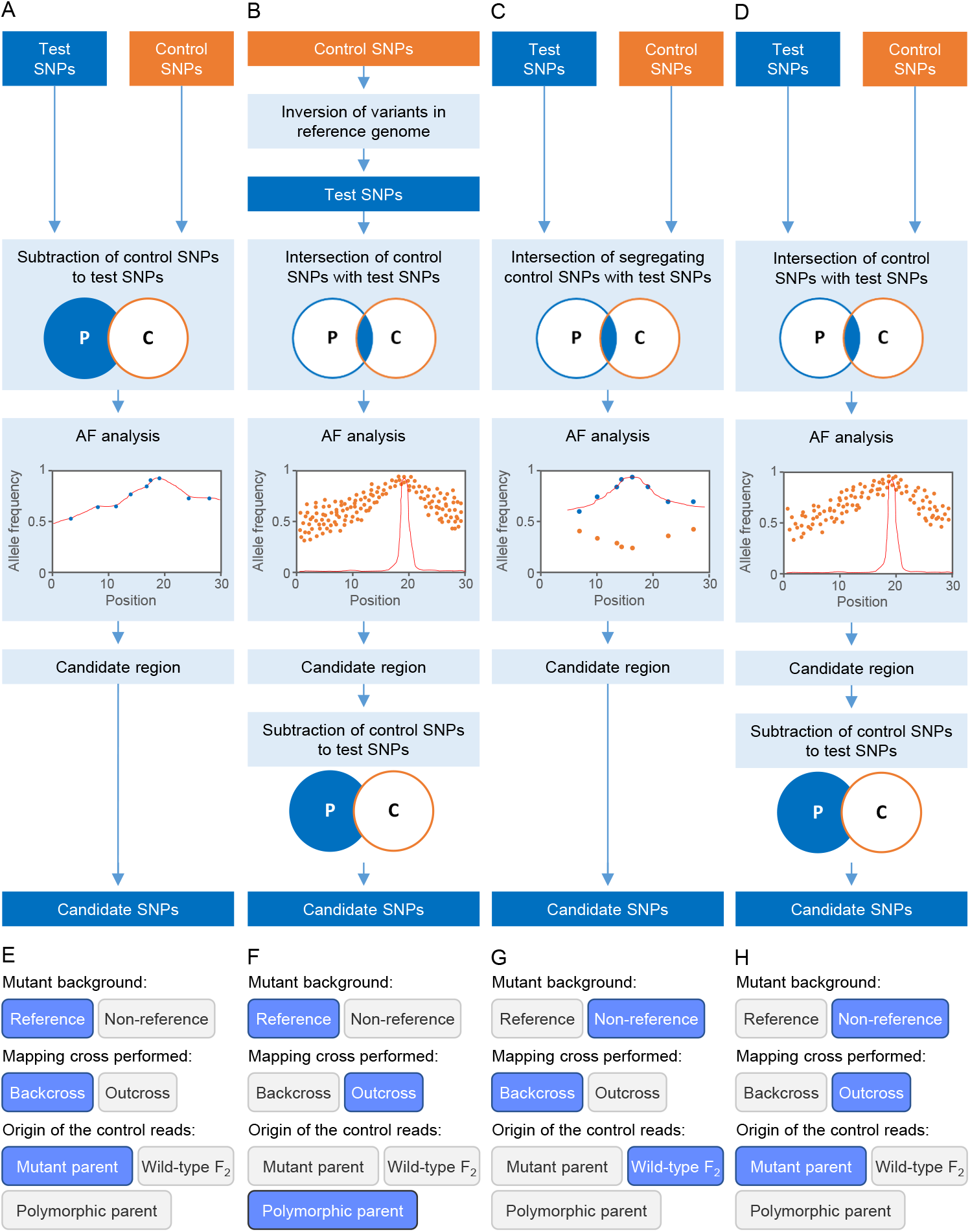
Some strategies for EMS-induced mutation mapping implemented in Easymap. (A–D) The input reads are processed into control and test SNP lists. The lists are contrasted in order to determine the SNPs that can be informative for mapping, which are subjected to an allelic frequency (AF) analysis to find the phenotypically selected genomic position. A candidate region is defined around the selected genomic position, and the potentially causal SNPs within the candidate region are collected as candidate SNPs. (A) For a mutant strain obtained in the reference genetic background, a backcross is performed to obtain the mapping population and the control sample used is the parental of the mutagenized line. (B) For a mutant obtained in the reference genetic background, an outcross is performed to obtain the mapping population, and the control sample is the polymorphic wild-type parent. (C) For a mutant obtained in a non-reference strain, a backcross is performed to obtain the mapping population, and the control sample used is a pool of phenotypically wild-type F_2_ individuals. (D) For a mutant obtained in a non-reference strain, an outcross is performed to obtain the mapping population, and the control sample is the parent of the mutagenized line. (E-H) Selection of the experimental design corresponding to panels A to D in the multiple-choice selectors of the graphic interface of Easymap.

**Table 1.**
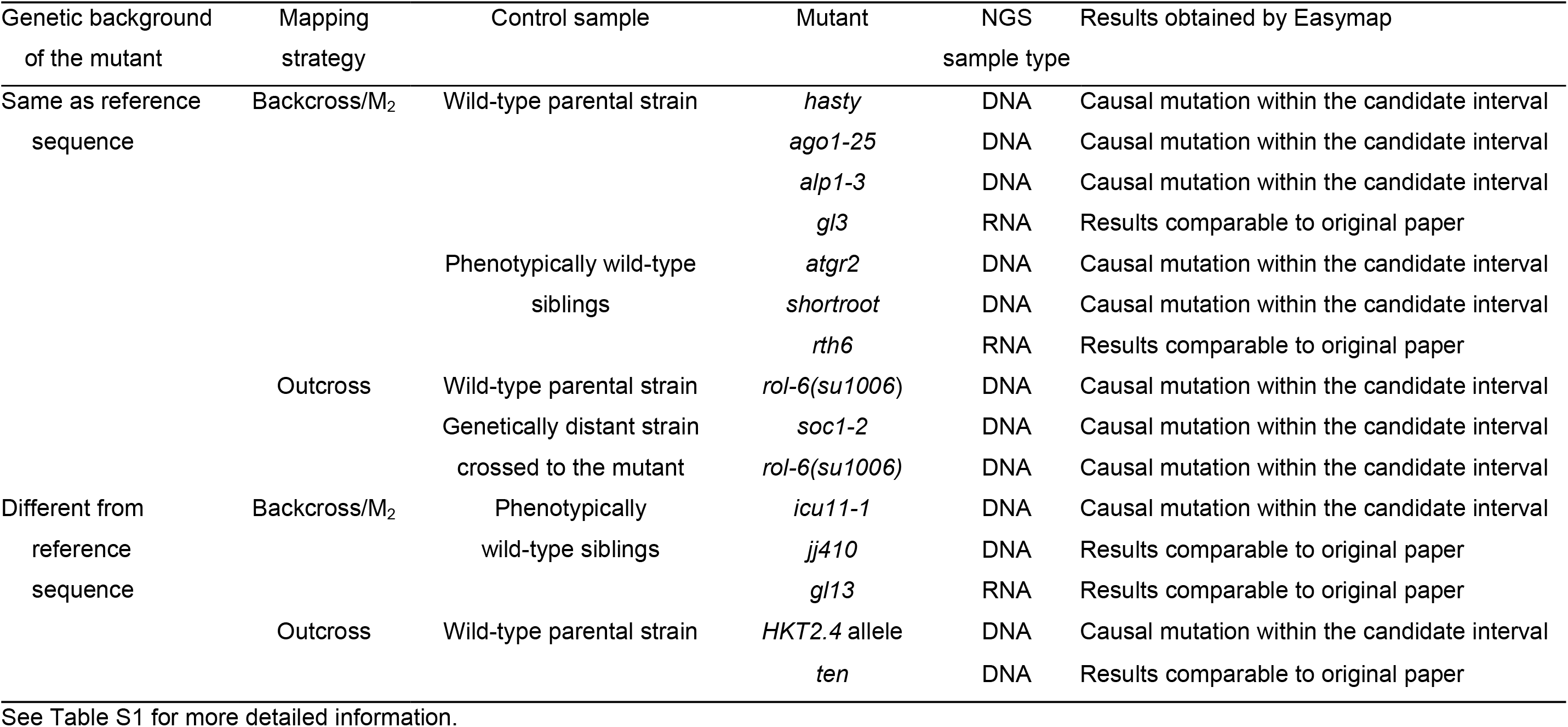
Validation of point-mutation mapping strategies using published, real experimental data

#### Workflow 2: Tagged sequence mapping

Easymap uses a tagged-sequence strategy to map the positions of large DNA insertions of known sequence (Figure 3). The user has to obtain paired-end (e.g., Illumina-like) or single-end (e.g., Ion Proton-like) NGS reads from a mutant carrying an insertion of partially or completely known sequence. Easymap can also use reads from multiple mutants pooled into a single DNA sample, in which case the minimum read depth recommended is 10× per mutant. The program finds reads that overlap the left and right junctions of a given insertion, as well as unpaired alignments neighboring the insertion site; then it uses them as probes against the whole genome sequence (Figure 3). If several hits accumulate around a locus, its physical position is reported as a putative insertion site.

**Figure 3.**
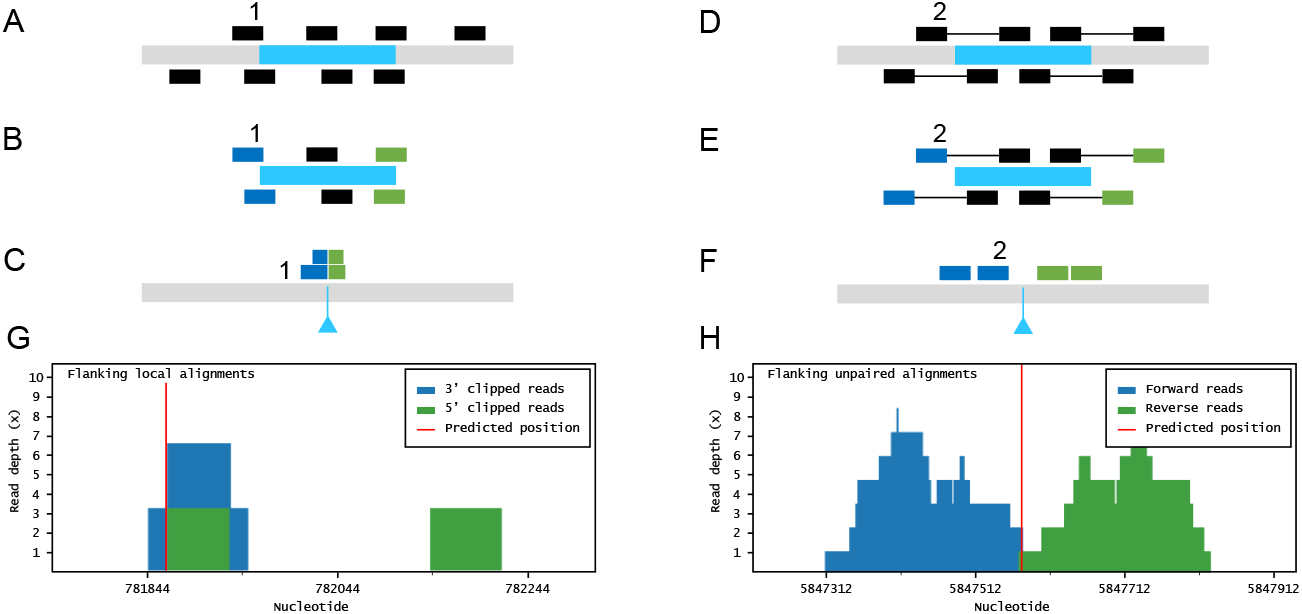
Large insertion mapping with Easymap. (A–C) Local alignment analysis. (A) The DNA insert appears in blue, over genomic DNA in grey. Individual reads are taken from the mutant genome. (B) The reads are aligned to the insertion sequence. Locally aligned reads (e.g., 1) are selected and sorted according to the end that is truncated (in blue and green). (C) The selected reads are aligned to the genomic reference sequence. The blue triangle indicates the position of the insertion in the mutant genome. (D–F) Paired-read analysis. (D) Paired reads are taken from the mutant genome. (E) The reads are aligned to the insertion sequence. Unaligned reads with aligned mates (e.g., 2) are selected and sorted according to their position in relation to the insertion (in blue and green). (F) The selected reads are aligned to the reference sequence, delimiting a candidate region for the insertion site. (G-H) Read depth histograms for examples of local alignment (G) and paired-read analyses (H). (G) False positive insertion, characterized by low overall read depths and disorganized data. (H) True positive insertion, characterized by high read depths and organized data.

If more than one insertional event is detected in a given mutant, the above experimental design and analysis workflow cannot discriminate the insertion causing the mutant phenotype. Such a mutant, however, can also be crossed and analyzed as described in Workflow 1. The mapping report from Workflow 2 includes a histogram that shows the distribution of the data supporting each putative insertion (Figure 1D). The user can inspect the histograms visually to easily discern false positives (disorganized clusters with very low accumulated read depths [RDs]) from genuine insertions (clusters of organized data with a high number of accumulated RDs).

#### Complementary tasks

An Easymap run automatically performs several essential tasks such as input data quality controls, including the verification of the FastQ encoding and quality, and assessment of the RD distribution for each sample. After each analysis, Easymap creates a comprehensive report containing high-resolution images and tabular data to assist the user in interpreting the mapping results (e.g., allele frequency plots for EMS-induced mutations, RD histograms for insertional mutations, and gene plots for each putatively damaged gene; Figure 1D), a prediction of the functional effect of the candidate mutations, the flanking sequences of each mutation, and the sequences of oligonucleotide primers to genotype such mutation.

NGS experiment simulations can be helpful for optimizing the design of effective mapping experiments (James et al., 2013; Wilson-Sánchez et al., 2019). Therefore, Easymap includes a built-in experiment simulator that allows the user to simulate NGS data in order to test different mapping designs and parameters.

### Assessment of Easymap performance with simulated and real data

We tested Easymap with tens of simulated datasets for each mapping strategy and analysis workflow supported by the program. This allowed us to hard-code appropriate values for analysis variables (e.g., SNP filtering thresholds) under different experimental conditions, saving the user from having to set complex parameters. However, to add more flexibility to the analysis, Workflow 1 allows the user to choose between two levels of stringency for SNP selection. We tested Easymap with data from real mapping-by-sequencing experiments; Tables 1 and 2 show the results that we obtained when analyzing reads from a range of previously published mutants. We reproduced previously published results, demonstrating the reliability of Easymap even under extreme conditions with average read depths as low as 5× (Obholzer et al., 2012; Wilson-Sánchez et al., 2014). The mapping reports for each of these experiments are available in our preview Easymap installation (http://atlas.umh.es/genetics/) and additional information about these experiments is provided in Table S1. Among the data used for testing the Easymap linkage analysis mapping workflows, we employed NGS reads obtained from mutants of *Arabidopis thaliana* (Morel et al., 2002; Hartwig et al., 2012; Rishmawi et al., 2014; Sun and Schneeberger, 2015; Wachsman et al., 2017; Mateo-Bonmatí et al., 2018), *Zea mays* (Liu et al., 2012; Li et al., 2013; Li et al., 2016a; Klein et al., 2018), *Caenorhabditis elegans* (Fay and Spencer, 2006) and *Danio rerio* (Obholzer et al., 2012), which included F_2_ (Hartwig et al., 2012; Obholzer et al., 2012; Rishmawi et al., 2014; Sun and Schneeberger, 2015; Klein et al., 2018; Mateo-Bonmatí et al., 2018), and M_2_/M_3_ (Wachsman et al., 2017) mapping populations obtained to identify recessive mutations, as well as dominant mutations mapped in F_2_ after an F_3_ screening (Fay and Spencer, 2006), and RNA-seq datasets (Liu et al., 2012; Li et al., 2013; Li et al., 2016a) (Tables 1 and S1). For tagged sequence mapping, we reproduced previous results for *Arabidopsis thaliana* (Wilson-Sánchez et al., 2014; Li et al., 2016b) and *Oryza sativa* (Yang et al., 2013) mutants; we also analyzed an unpublished dataset obtained in our laboratory (Tables 2 and S1).

**Table 2.**
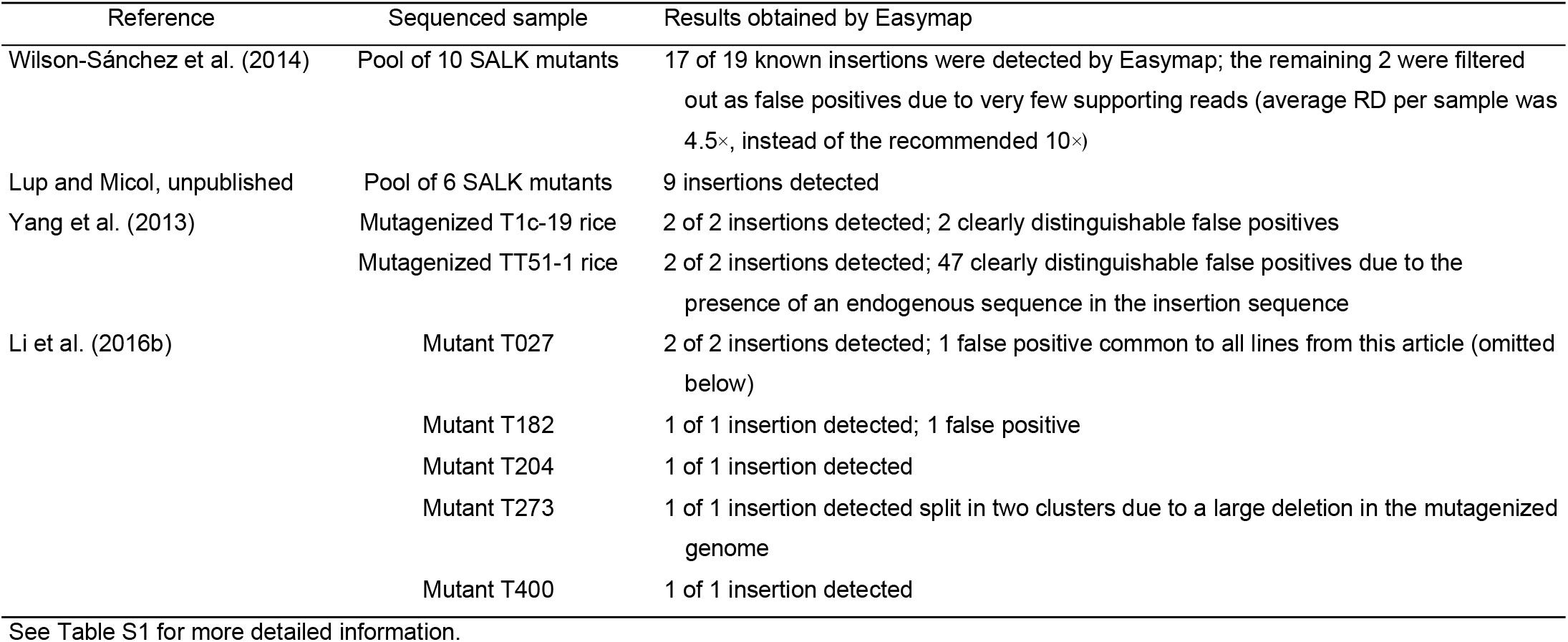
Validation of large-insertion mapping strategies with real experimental data

| Reference | Sequenced sample | Results obtained by Easymap |
| --- | --- | --- |
| Wilson-Sánchez et al. (2014) | Pool of 10 SALK mutants | 17 of 19 known insertions were detected by Easymap; the remaining 2 were filtered out as false positives due to very few supporting reads (average RD per sample was 4.5×, instead of the recommended 10×) |
| Lup and Micol, unpublished | Pool of 6 SALK mutants | 9 insertions detected |
| Yang et al. (2013) | Mutagenized T1c-19 rice | 2 of 2 insertions detected; 2 clearly distinguishable false positives |
|  | Mutagenized TT51-1 rice | 2 of 2 insertions detected; 47 clearly distinguishable false positives due to the presence of an endogenous sequence in the insertion sequence |
| Li et al. (2016b) | Mutant T027 | 2 of 2 insertions detected; 1 false positive common to all lines from this article (omitted below) |
|  | Mutant T182 | 1 of 1 insertion detected; 1 false positive |
|  | Mutant T204 | 1 of 1 insertion detected |
|  | Mutant T273 | 1 of 1 insertion detected split in two clusters due to a large deletion in the mutagenized genome |
|  | Mutant T400 | 1 of 1 insertion detected |

See Table S1 for more detailed information.

### Easymap architecture

We developed Easymap for UNIX-based operating systems since high-performance machines most commonly run Linux, and several tasks in Easymap are performed by third-party software that has already been extensively tested and used in Linux. These open-source, publicly available programs obtained by previous authors are listed in Table 4.

**Table 3.** Experimental designs supported by different open-source programs used for mapping-by-sequencing

| Mutations under study | Mapping design | Control sample | Software |  |  |  |  |
| --- | --- | --- | --- | --- | --- | --- | --- |
|  |  |  | SHOREmap<sup>1</sup> | SIMPLE<sup>3</sup> | CandiSNP<sup>4</sup> | Jitterbug<sup>5</sup> | Easymap<sup>7</sup> |
|  |  |  | ARTmap<sup>2</sup> |  |  | ITIS<sup>6</sup> |  |
| Point mutations | Backcross | Parental line | D |  | D |  | D/R |
|  |  | Phenotypically wild-type F<sub>2</sub> or M<sub>2</sub> |  | D |  |  | D/R |
|  | Outcross | Parental line | D |  |  |  | D/R |
|  |  | Phenotypically wild-type F<sub>2</sub> or M<sub>2</sub> |  | D |  |  | D/R |
| Large insertions | – | – |  |  |  | D | D |

The capabilities of some current mapping tools are compared, each character (D, with DNA-seq data; R, with RNA-seq data) representing an experimental design supported by the software. <sup>1</sup>Sun and Schneeberger (2015). <sup>2</sup>Javorka et al. (2019). <sup>3</sup>Wachsman et al. (2017).<sup>4</sup>Etherington et al. (2014). <sup>5</sup>Hénaff et al. (2015). <sup>6</sup>Jiang et al. (2015). <sup>7</sup>This work.

**Table 4.**
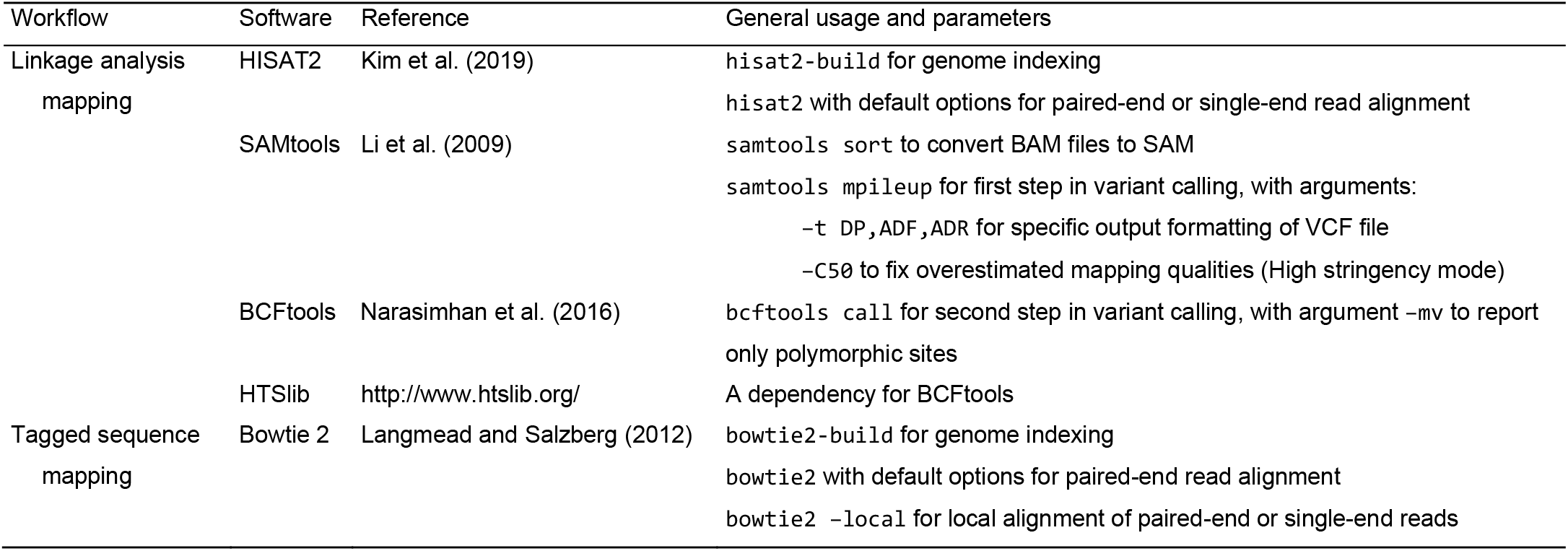
Third party software packages included in Easymap

| Workflow | Software | Reference | General usage and parameters |
| --- | --- | --- | --- |
| Linkage analysis<br>mapping | HISAT2 | Kim et al. (2019) | hisat2-build for genome indexing<br>hisat2 with default options for paired-end or single-end read alignment |
|  | SAMtools | Li et al. (2009) | samtools sort to convert BAM files to SAM<br>samtools mpileup for first step in variant calling, with arguments:<br>-t DP,ADF,ADR for specific output formatting of VCF file<br>-C50 to fix overestimated mapping qualities (High stringency mode) |
|  | BCFtools | Narasimhan et al. (2016) | bcftools call for second step in variant calling, with argument -mv to report only polymorphic sites |
|  | HTSlib | http://www.htslib.org/ | A dependency for BCFtools |
| Tagged sequence<br>mapping | Bowtie 2 | Langmead and Salzberg (2012) | bowtie2-build for genome indexing<br>bowtie2 with default options for paired-end read alignment<br>bowtie2 -local for local alignment of paired-end or single-end reads |

Easymap comprises a software stack consisting of a controller layer, a workflow layer (representing linkage analysis mapping, tagged sequence mapping, and common processes), and a tasks layer (representing custom and third-party programs). The controller exposes a simple application programming interface (API) with which the web and command line scripts interact. This allows Easymap to be installed either locally or remotely while offering simultaneous command-line and graphical interface access (Figures S1 and S2).

To simplify Easymap installation, a single script compiles and/or installs all required software (Python2, Python Imaging Library, Virtualenv, Bowtie2, HISAT2, HTSlib, SAMtools, and BCFtools; Table 4) within the Easymap directory. All third-party software is included within the Easymap package so that no dependencies are required. For installations in shared environments, usage (memory and number of concurrent jobs) can be limited by the system administrator through a simple configuration file. The installation script sets up a dedicated HTTP server to run as a background process using the port chosen by the user. Easymap implements chunked file transfers for reliable HTTP transfer of large read files. Further installation setups and usage indications can be found in the Easymap documentation, including indications for usage within the Windows 10 operating system (File S1).

## DISCUSSION

Hundreds of point and insertional *Arabidopsis thaliana* mutants have been isolated in our laboratory over almost 30 years. After the advent of NGS, we generated NGS datasets to identify the mutations causing the mutant phenotypes of interest and analyzed these datasets using different available software tools. We realized that most tools intended for mapping-by-sequencing are not easily manageable by researchers without a background in bioinformatics. We attempted to identify the main accessibility issues of such tools and developed Easymap, a program for mapping-by-sequencing that can be used reliably by as many researchers as possible, irrespective of their computer skills, and in as many experimental designs as possible.

The main accessibility features of Easymap are the following: it is free and open source; a single command installs the software and launches the server for the graphical interface; it is easy to use, as it has a graphical interface and workflows that smoothly convert raw data into comprehensive yet simple reports; it is polyvalent, since it can be used for a wide variety of experimental setups (Table 3); and it is flexible, since it can be installed locally or remotely (on a server) while maintaining its graphical interface (Figures S1 and S2).

The implementation of the HISAT2 aligner allows the use of RNA-seq data for mutation mapping in large genomes, making Easymap the first program of its class to allow perfoming mapping by sequencing with large genomes for which whole genome sequencing may not be affordable. Easymap proved to be reliable under a wide variety of experimental designs, in five different plant (*Arabidopsis thaliana*, *Zea mays* and *Oryza sativa*) and animal (*Danio rerio* and *Caenorhabditis elegans*) species and a total of 28 experiments showing unprecedented versatility and adaptability to the input data.

In conclusion, here we introduce Easymap, a novel analysis tool for mapping-by-sequencing of large insertions and point mutations, which has been designed to accommodate all potential users. Easymap features a web-based graphic interface, a simple installation script, robust mapping analyses for several experimental designs and thorough user-oriented mapping reports. A preview instance of Easymap is available at http://atlas.umh.es/genetics, where we also offer a quickstart installation guide (File S2). The easymap source code is available for download at https://github.com/MicolLab/easymap and http://genetics.edu.umh.es/resources/easymap.

## METHODS

### Programming languages and utilities

Easymap was designed as a modular program, so that each module can be used and modified independently. Modules include custom Python2 scripts and third-party software packages (Table 4). Modules are run sequentially by different Bash scripts, or workflows, attending to the user preferences as defined in the web interface or the command line interface.

Python source and libraries are installed within a Virtualenv virtual environment so that previous software installations are not disturbed. The Easymap server is launched using the Python2 CGIHTTPServer function to set up the web interface. The Pillow imaging library is used for the generation of the graphic output.

### Testing

We tested Easymap in several operating systems, including different distributions of Linux such as Ubuntu, Fedora, Red Hat, and AMI. Easymap can also run within the Ubuntu apps available in the Windows 10 Microsoft Store. Easymap runs on regular desktop computers and high-performance machines, in local machines and remote instances (e.g., the Amazon Elastic Compute Cloud service), and also in virtual machines running UNIX-based OS within Windows OS. Appendix D of the Easymap user manual (File S1) includes detailed information for different installation setups.

## ACKNOWLEDGEMENTS

This work was supported by grants from the Ministerio de Ciencia e Innovación of Spain (PGC2018-093445-B-I00 [MCI/AEI/FEDER, UE]) and the Generalitat Valenciana (PROMETEO/2019/117) to JLM.

## SUPPLEMENTAL INFORMATION

**File S1. Easymap documentation.**

**File S2. Easymap quickstart guide.**

**File S3. Supplemental figures.**

**Table S1. Results obtained in the validation of Easymap using multiple NGS datasets.**

## Author contributions

J.L.M. obtained funding, provided resources and supervised this work. D.W., S.D.L, and J.L.M. conceived and designed the program; D.W. and S.D.L. developed the program, S.A. contributed a number of Python scripts; S.D.L. tested the software with real datasets. S.D.L., D.W. and J.L.M wrote the article.

## Abbreviations

NGS: next-generation sequencing
EMS: ethyl methanesulfonate
RD: read depth
CV: coverage
SNP: single-nucleotide polymorphism
AF: allele frequency

